# Missing the forest because of the trees: Slower alternations during binocular rivalry are associated with lower levels of visual detail during ongoing thought

**DOI:** 10.1101/2019.12.31.891853

**Authors:** Nerissa Siu Ping Ho, Daniel Baker, Theo Karapanagiotidis, Paul Seli, Hao Ting Wang, Robert Leech, Boris Bernhardt, Daniel Margulies, Elizabeth Jefferies, Jonathan Smallwood

## Abstract

Conscious awareness of the world fluctuates, either through variation in the quality with which we perceive the environment, or, when attention switches to information generated using imagination rather than the external environment. Our study combined individual differences in experience sampling, psychophysical reports of perception, and neuroimaging descriptions of structural connectivity, to better understand these changes in conscious awareness. In particular, we (a) examined if aspects of ongoing thought, as measured by multi-dimensional experience sampling during a sustained attention task, are associated with the white matter fibre organization of the cortex as reflected by their relative degree of anisotropic diffusion, and (b) whether these neuro-cognitive descriptions of ongoing experience are related to a more constrained measure of visual consciousness provided by the analysis of bistable perception during binocular rivalry. Individuals with greater fractional anisotropy (FA) in right hemisphere white matter regions involving the inferior fronto-occipital fasciculus, the superior longitudinal fasciculus and the cortico-spinal tract, described their ongoing thoughts as lacking external details. Subsequent analysis indicated that the combination of low FA in these right hemisphere regions, with reports of high level external details, was associated with the shortest periods of dominance during binocular rivalry. Since variation in binocular rivalry reflects differences between bottom-up and top-down influences on vision, our study suggests that reports of ongoing thoughts with vivid external details may occur when conscious precedence is given to bottom-up representation of perceptual information and that this may partly be rooted in the white matter fibre organization of the cortex.

## Introduction

Conscious experience varies from moment to moment, and studies using experience sampling suggest that these dissociations can take multiple forms. Sometimes our thoughts become focused on personal experiences rather than events in the external environment, or any task being performed (Seli et al., 2018; Smallwood and Schooler, 2006, 2015). Research suggests states of off-task thought are linked to systems important for attentional focus, including both dorsal and ventral attention networks (Hasenkamp et al., 2012; Turnbull et al., 2019a; Turnbull et al., 2019b). Experience can also fluctuate in the level of details with which events in the external environment are processed. Recent studies using experience sampling highlight situations when external events are experienced in a detailed manner depend on the functioning of the default mode network (DMN, Sormaz et al., 2018), in particular the posterior cingulate and para-hippocampus (Ho et al., 2019; Murphy et al., 2019; Turnbull et al., 2019b). Although a link between the DMN and externally focused experience is surprising given the widely held assumption of this network as being limited to internally focused, task-negative experiences, it is nonetheless consistent with evidence of a role when individuals perform tasks with high levels of efficiency such as when they “in the zone” (Esterman et al., 2012; Kucyi et al., 2016) or “on autopilot” (Vatansever et al., 2017). Understanding the neural mechanisms underlying different patterns of experience, is therefore, not only important for contemporary accounts of ongoing conscious thought (Smallwood and Schooler, 2015), but, may also be important for appropriately characterising the function of different large scale neural networks.

The current study, therefore, aimed to understand the role that top-down visual processes play in different dissociations between experience and events in the immediate environment. Traditionally, research into conscious experience has emphasised that it is possible to understand the relationship between subjective awareness and the immediate sensory context using situations of bistable perception (such as the Necker Cube, or the phenomenon of binocular rivalry) because in this context awareness can change without a concomitant change in sensory input (Crick, 1996). Situations of bistable perception, therefore, provide relatively unambiguous indices of the top down influence on vision because they discard low-level processes that contribute to the process of perception (e.g. sensory transduction) and yet may not be directly related to awareness. In this context, when one percept dominates during bistable perception this is assumed to reflect top-down influences on vision. Consistent with the assumption that bistable perception depends on the balance between top-down and bottom-up influences on vision, neuroimaging studies suggest that rivalry depends on both processes taking place in visual regions (Tong et al., 2006) as well as higher-order brain regions (Baker et al., 2015; Knapen et al., 2011). Importantly, both default mode and attention systems are important in binocular rivalry: Disruptions to regions of the default mode network tend to slow down perceptual alterations during bistable perception (Carmel et al., 2010; Kanai et al., 2011) while disruptions to the dorsal attention network shorten perceptual alterations (Kanai et al., 2011).

Our study sought to extend our understanding of naturally occurring changes in ongoing experience, by linking them to both changes in the structural organisation of the cortex, as well as to indices of the top-down influence on vision as estimated from binocular rivalry alternations. Specifically, we analysed data from a large cohort of individuals for whom we had extensively described the contents of their ongoing experience in a laboratory task (for prior publications see Ho et al., 2019; Sormaz et al., 2018; Turnbull et al., 2019a; Turnbull et al., 2019b; Wang et al., 2018) and for whom we also acquired measures of binocular rivalry using a similar paradigm to our prior study (see Baker et al., 2015). These individuals also had measures of structural connectivity provided by diffusion tensor imaging (DTI), which has highlighted neural processes linked to both binocular rivalry (Genç et al., 2011) and to patterns of ongoing thought in a prior study (Karapanagiotidis et al., 2017). In that study we found a region of right lateralized white matter, that had greater fractional anisotropy (FA) for individuals who tended to neglect the external environment by imagining events in the past or future instead of those in the here and now. Our current study aimed to replicate the association between ongoing experience and white matter architecture of the right hemisphere in a new set of participants and then explore whether this association was related to the relative balance between top-down and bottom-up influences on vision during binocular rivalry, as indexed by an individual’s reported experience during binocular rivalry.

The left hand panel in Figure 1 describes the tasks we used to measure ongoing experience during a sustained attention task, as well as binocular rivalry. Using these data, in combination with metrics of white matter architecture provided by DTI, we set out to answer two questions: (1) Are individual differences in pattern of one’s ongoing thoughts reflected in the structure of cortical white matter? and (2) Whether these neuro-cognitive descriptors for the patterns of ongoing thoughts are related to more precise descriptions of perceptual experience as assessed by measures of binocular rivalry?

**Figure 1.**
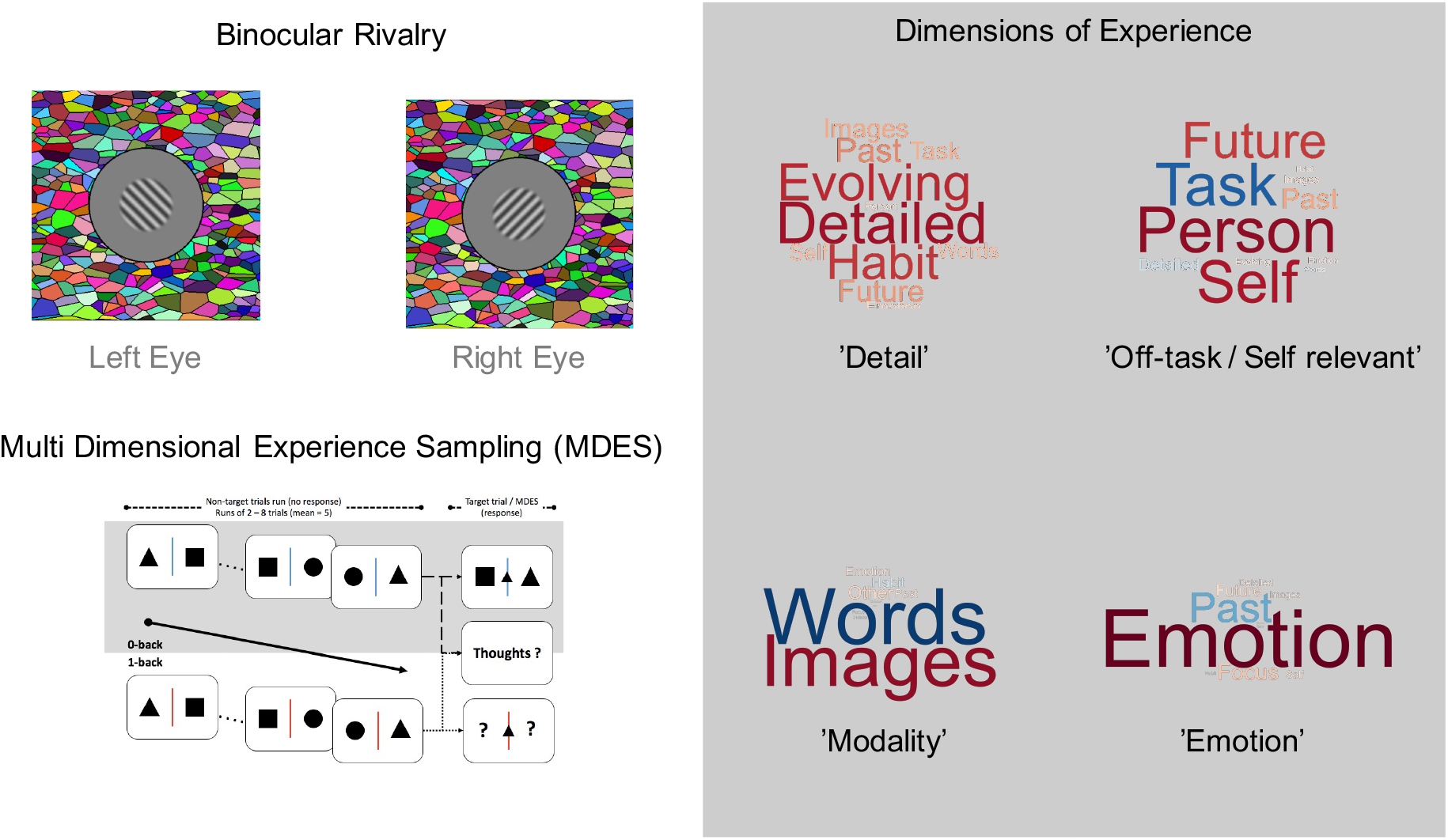
Experimental Protocol. Participants participated in laboratory sessions in which we measured conscious experience through the estimation of binocular rivalry during bistable perception (top left) and also using experience sampling while they performed a simple non-demanding cognitive task (bottom left). Application of principal components analysis (PCA) to the Multi-Dimensional Experience Sampling (MDES) revealed four components which are displayed in the form of word clouds on the right hand side panel. The colour and size of the words indicate the loadings of each question (font size = strength of relationship and colour = direction). The labels we used to describe these components in the paper are presented in quotations.

## Methods

### Participants

150 healthy, right-handed, native English speakers, with normal or corrected-to-normal vision and no history of psychiatric or neurological illness (Mean Age = 20.19 and 92 were females) participated in the study. All participants provided written informed consent approved by the Department of Psychology and York Neuroimaging Centre (YNIC), University of York ethics committees, and were debriefed after completion of the study. They were either paid or given course credit for their participation.

### Procedures

Participants arrived at YNIC where we acquired brain images including T1 Weighted MRI, resting state MRI and diffusion tensor imaging DTI. On subsequent days, participants took part in a comprehensive set of behavioural assessments that captured different aspects of cognition including both the experiential assessment task and other experimental tasks (including binocular rivalry). These tasks were completed over three sessions on different days, with the order of sessions counterbalanced across participants. The task in which ongoing experience was measured always took place at the start of these laboratory sessions.

### Experience Sampling

We measured ongoing cognition in a paradigm that manipulates memory load by using alternating blocks of 0-back (low-load) and 1-back (high-load) conditions, with the initial block counter-balanced across individuals (see Turnbull et al., 2019b for a complete description of this task). Multi-dimension Experience Sampling (MDES) was used to measure the contents of on-going thought. On each occasion when probed with ‘Thought ?’ (see bottom left panel of Figure 1), participants reported their thoughts by responding to one of the 13 questions presented in Supplementary Table 1. Participants always rated their task focus first and then described their thoughts at the moment before the probe on a further 12 dimensions.

### Binocular Rivalry

We showed rivalling stimuli to participants for four trials of 120 s in duration and asked them to report their percepts using a computer mouse. The stimulus consisted of oblique gratings (1c/deg, 50% contrast, ±45 deg, 6 deg in diameter, smoothed by a raised cosine envelope) shown to opposite eyes (see top left panel of Figure 1). All stimuli were presented on a gamma-corrected Iiyama VisionMaster Pro 510 CRT monitor with a mean luminance of 32 cd/m^2^, and were viewed through a mirror stereoscope to permit presentation of different images to the left and right eyes. The stimuli were surrounded by a dark ring, and a binocular Voronoi texture to promote binocular vergence and fusion (Baker and Graf, 2009). Participants held down one mouse button when they perceived a particular percept (e.g. a left-oblique grating) and the other when they perceived the alternative (e.g. a right-oblique grating). If they simultaneously perceived both, or experienced a mixed percept, they held down both buttons. We counterbalanced the orientations of the rivalling stimuli between the eyes on alternate trials.

### Diffusion Tensor Imaging

The DTI scan lasted 13 minutes. A single-shot pulsed gradient spin-echo echo-planar imaging sequence was used with the following parameters: b = 1000 s/mm^2^, 45 directions, 7 T2-weighted EPI baseline scans, 59 slices, FOV = 192 × 192 mm^2^, TR = 15 s, TE = 86 ms (minimum full), voxel size = 2 × 2 × 2 mm^3^, matrix = 96 × 96. DTI data preprocessing steps involved eddy-current distortion correction and motion correction using FDT v3.0, part of FSL (Smith et al., 2004). FA was calculated by fitting a tensor model at each voxel of the preprocessed DTI data and the resulting images were brain-extracted using BET (Smith, 2002). Voxel-wise FA maps were analysed using Tract-Based Spatial Statistics (TBSS) (Smith et al., 2006). After subjects’ FA data were non-linearly aligned to FMRIB58_FA standard space, they were transformed to the mean space of these subjects and then affine transformed to the 1 mm MNI152 space. Next, the mean of all FA image was created and thinned to create a mean FA skeleton which represents the centres of all tracts common to the group.

The skeletonised FA images were then fed into voxelwise statistics, using FSL’s Randomize (a nonparametric permutation inference tool). Using a generalised linear model (GLM), the measured FA values across the skeleton were regressed with the experience sampling results, while age and gender were included as nuisance covariates. T-statistic maps for contrasts of interest were calculated with 5000 permutations (Nichols and Holmes, 2002). Resulting maps were thresholded at a Family-Wise Error (FWE) corrected *p*-value of .05 using Threshold-Free Cluster Enhancement (TFCE) (Smith and Nichols, 2009).

Probabilistic diffusion models were also fitted using BEDPOSTX (Behrens et al., 2003) with 2 fibres modelled per voxel and 1000 iterations. Probabilistic tractography was performed using PROBTRACKX (Behrens et al., 2007) to reconstruct fibres passing through the ROI resulted from the above GLM analysis if high degree of cross fibres existed (see below Results: Associations with white matter fibre organization). Tractography was performed in native diffusion space by transforming the ROI as seed masks from standard space into diffusion space using the inverse of the nonlinear registration calculated in the TBSS pipeline. We used standard parameters (5000 samples/voxel, curvature threshold 0.2, step length 0.5 mm, samples terminated after 2000 steps or when they reached the surface as defined by a 40% probabilistic whole-brain WM mask). The connectivity maps of each individual were thresholded at 1% of total samples, mapped to standard space using nonlinear registration and concatenated into a single 4D file.

## Results

### Categorising experience

#### Binocular rivalry

Two metrics were calculated using data from the bistable perception session. The first was the mean duration (in seconds) of each period where one stimulus continuously dominated experience (‘Dominance Duration’). Mean dominance duration shows robust and stable individual differences (Pettigrew and Miller, 1998), which have previously been shown to be associated with connectivity between regions of parietal cortex (Baker et al., 2015), as well as the concentration of inhibitory neurotransmitters (GABA) in visual regions of the brain (van Loon et al., 2013). Dominance durations are also affected by various personality types (Antinori et al., 2017a; Antinori et al., 2017b) and clinical conditions including autism (Robertson et al., 2013), bipolar disorder (Miller et al., 2003; Pettigrew and Miller, 1998) and schizophrenia (Xiao et al., 2018; Ye et al., 2019). The second metric was the time when neither percept dominated experience (‘Mixed’). Mixed percepts occur at transitions between states of full dominance, and involve a network of frontal and parietal brain areas, particularly in the right hemisphere (Knapen et al., 2011). Measures of these metrics were then transformed into z-scores, with outliers (> 2.5, and based on visualisation of boxplot generated in SPSS 25) being replaced with mean values (outliers: ‘Dominance Duration’ = 23, ‘Mixed’ = 11). We found no correlations on the scores of these two metrics (*r* = .02, *p* < .9).

#### Experience sampling

In our analysis, we used the decomposition reported in Sormaz colleagues (2018, see original paper for complete details). In brief, principal components analysis (PCA) was applied to MDES data at the trial level as is standardized in our other works (for examples see Konishi et al., 2017; Smallwood et al., 2016). This produced four components: (1) ‘Detail’, reflecting patterns of detailed visual task related experience, (2) ‘Off-task thought’, dissociating on task thought from episodic self-relevant thought, (3) ‘Modality’, distinguishing thoughts related to images or words and (4) ‘Emotion’ describing the affective tone of experiences. These components are presented in the form of word clouds in the right hand panel of Figure 1.

### Associations with white matter fibre organization

Our first analysis examined associations between white matter connectivity and the patterns of ongoing thought identified using MDES data. First, we identified regions of white matter where FA varied across individuals with the patterns of thought that they reported, by conducting a multiple regression in which skeleton wide FA maps for each individual were the dependent variable. Each individual’s score for each of the experiential dimensions identified through PCA were explanatory variables. Age and gender were included as nuisance covariates. Significant negative associations between FA and detailed thoughts were identified, and regions showing this relationship are presented in red in Figure 2.

**Figure 2.**
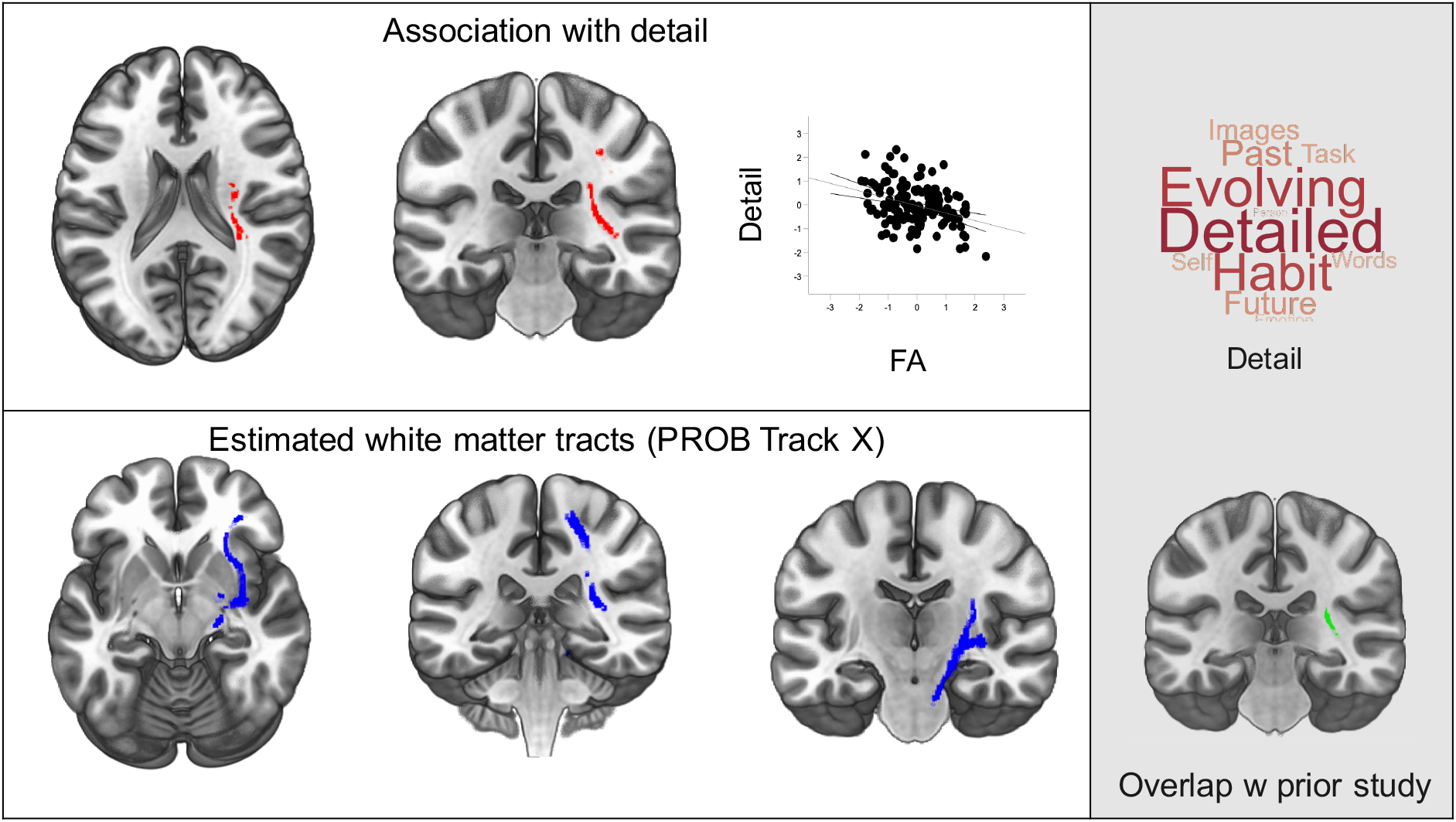
White matter tracts associated with less detailed external experience. Regions shown in red represent areas where FA was higher for individuals whose thoughts lacked vivid detail. In the scatter plot each point is an individual (top left). Estimates of the most probable white matter tracts (bottom left). Brain maps were corrected for family-wise error using TFCE (Threshold-Free Cluster Enhancement) thresholding method (*p* < .05 FWE-corrected). The grey panel on the right shows the pattern of responses that reflect high levels of external detail (top right) and the overlap between the result in the current study with our prior study that forms the basis of the PROBTRAKX analysis (bottom right).

Next, we examined the relationship between the current result and those from our prior study (Karapanagiotidis et al., 2017). In our prior work using different participants, we identified a set of right lateralized tracts with greater FA for individuals reporting more mental time travel. Comparison of the two FWE-corrected maps indicated an area of overlap (see bottom right panel in Figure 2). Our prior study found higher FA linked to experiences characterised by thoughts focused on the self-generation of thoughts about the past and the future, while the current results highlighted higher FA was linked to less detailed assessments of the here and now. Together, therefore, these two results provide converging evidence that right lateralized white matter tracts are important for differences in internal versus external focus of attention. As this region of overlap has a high degree of crossing fibres, we used ProbTrackX to estimate the white matter bundles to which this was most likely to be related (See Methods, bottom left panel of Figure 2). It can be seen that the results of this process highlight multiple large fibre bundles including the inferior occipital-frontal (IFOF) and the cortico-spinal tract (CST), and the superior longitudinal fasciculus (SLF).

### Associations between different features of conscious experience

Having documented associations with white matter structure and ongoing thought, we next examined: (a) Are there patterns of ongoing experience identified by MDES that are related to the nature of experience as determined via binocular rivalry? and (b) If so, whether these associations are related to associated white matter differences in brain structure. Table 1 shows the zero order relationships between this set of variables. It can be seen that a weak correlation is present between ‘Detail’ thoughts and ‘Difference’ (the difference between z-scores of mean dominance duration and the proportion of mixed percepts). This indicates that individuals with more detailed thoughts experience a combination of longer periods of dominance and fewer mixed percepts.

**Table 1.** Simple correlations between z-scored measures of ongoing experience (represented as the rows) and z-scored metrics of bistable perception (represented in the columns). The difference column describes the difference between z-scores of mean dominance duration and the proportion of mixed percepts. *r* = correlation value; *p* = *p*-value.

|  |  | **‘Dominant’** | **‘Mixed’** | **Difference** |
| --- | --- | --- | --- | --- |
| **Detail** | *r* | -0.12 | 0.13 | 0.18 |
|  | *p* | 0.13 | 0.13 | 0.03* |
| **Off Task** | *r* | -0.07 | 0.05 | 0.08 |
|  | *p* | 0.42 | 0.57 | 0.34 |
| **Modality** | *r* | -0.07 | 0.01 | 0.05 |
|  | *p* | 0.41 | 0.94 | 0.54 |
| **Modality** | *r* | -0.07 | 0.14 | 0.15 |
|  | *p* | 0.42 | 0.09 | 0.07 |

To formally understand the relationship between different patterns of thought, their observed associations with white matter architecture, and the estimates of experience provided by binocular rivalry, we conducted a Multivariate Analysis Of Co-Variance (MANCOVA). In this analysis, mean dominance duration and the proportion of mixed percepts were the dependent variables. Individual scores on each PCA dimension, as well as the DTI correlate of detailed experience (that is, the mean FA for the white matter region that is correlated with detailed experience), were explanatory variables, while age and gender were included as nuisance covariates. We modelled the main effect of each explanatory variable, as well as the interaction between ‘Detail’ and its white matter correlate. We found a significant interaction between ‘Detail’ and its association with white matter connectivity (*F*(2, 140) = 4.8, *p* = .012), reflecting differences in mean dominance duration (*F*(1, 149) = 7.22, *p* = .008). Plotting the association between dominance duration and FA separately for individuals with high and low Detail (using median split method) revealed the shortest dominance durations were observed in individuals with high levels of Detail and the lowest FA (see left hand panel of Figure 3). Assuming that longer states of rivalry are linked to more top-down influence over vision, then our results suggest that experiencing the external world with high levels of detail is linked to relatively greater bottom-up influences on visual experience.

**Figure 3.**
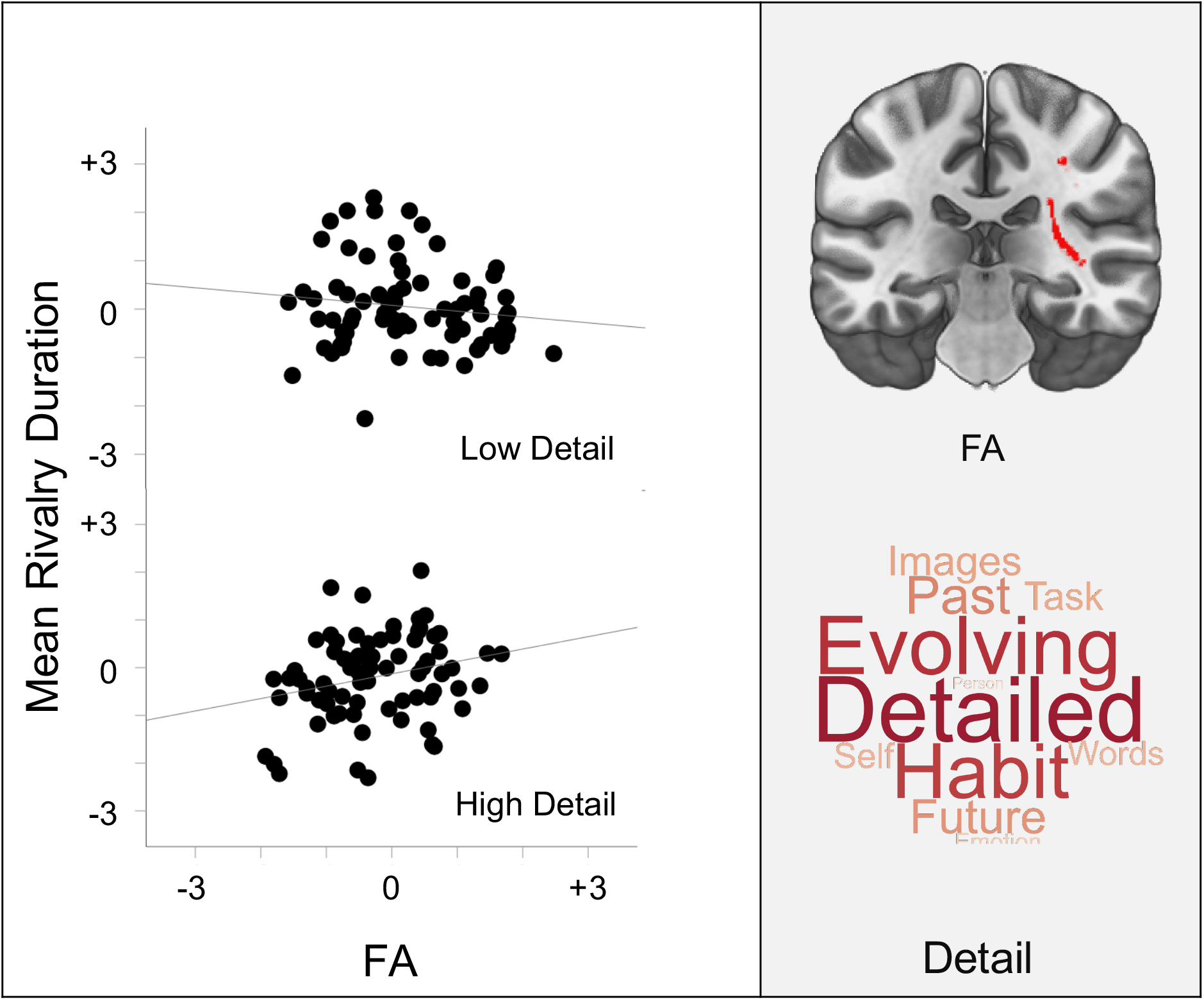
Association between individual variations in external detail in ongoing experience derived through experience sampling, the associated white matter architecture and visual experience determined via binocular rivalry. Scatter plots show that the longest periods of binocular rivalry dominance were experienced by participants with a combination of higher FA and reports of high external details in ongoing cognition (bottom left). For the purpose of display the sample was split at the median value on their reported levels of Detail.

## Discussion

Our study set out to leverage the methods of experience sampling, psychophysics and structural brain imaging to better understand shifts in the qualities of conscious experience. We found a correlation between individual differences in estimates of the integrity of cortical white matter in the right hemisphere and the level of detail with which external events were experienced. Notably, the pattern of right lateralized white matter tracts that had a greater integrity for less detailed experiences of the moment in these analyses, partially overlapped with our prior analysis of a different sample highlighting greater FA for individuals with a greater focus <u>away</u> from the moment to other times and places (Karapanagiotidis et al., 2017). Given that external focus is reduced during periods of self-generated imaginative thought (Kam et al., 2011; Kam and Handy, 2013), these two results help establish the importance of a right lateralised network of white matter tracts in determining self-generated experience.

Using psychophysics we found that individuals who had the longest periods of dominance during bi-stable perception, reported the high level of external details during sustained attention and had the greatest estimates of white matter in these right lateralized regions. It is usually assumed that during rivalry, top-down processes stabilise one potential interpretation of visual input, and so shorter time during rivalry is related to a bias towards bottom-up influences. Based on our data, our participants reports of detailed experience during sustained attention may emerge because of a conscious emphasis on bottom-up influences derived from sensory input that is in part constrained by the white matter architecture of the cortex.

In this context it is interesting to note that our prior studies found that neural signals within the DMN (traditionally assumed to be linked to internal states) encode patterns of detailed thoughts focused on a working memory task (Sormaz et al., 2018), while Turnbull and colleagues (2019b) found, in the same cohort as we report here, that at rest greater functional connectivity between the DMN and with visual cortex, predicted more detail experiences in the laboratory. Our recent work on the macrostructural organisation of the cortex suggests that the DMN is functionally and spatially isolated from sensory and motor systems (Margulies et al., 2016). Perhaps the apparent contribution of the DMN to greater focus on external information with greater detail (Sormaz et al., 2018; Turnbull et al., 2019a; Turnbull et al., 2019b), or, during efficient task performance (e.g. Esterman et al., 2012; Vatansever et al., 2017), arises when there is a particularly strong representation of bottom up sensory signals in the DMN. Intriguingly, recent retinotopic mapping studies have identified that regions of the DMN can show patterns of selective deactivation as a function of the location of a visual stimulus (Szinte et al., 2019).

The fibre bundles identified through probabilistic tractography in our study, therefore, may offer a possible window into how the DMN can contribute to modes of operation that have both internal and external features. We found a region in both the current study and in our prior analysis using independent data (Karapanagiotidis et al., 2017) that was at the overlap of three major white matter fibre bundles. The cortico-spinal tract (CST) originates in regions of sensory and motor cortex and is thought to be important for voluntary behaviour, as well as modulation of sensory information (Kolb and Whishaw, 2009). The superior longitudinal fasciculus (SLF) is a major white matter pathway that connects the frontal, parietal, temporal and occipital lobes. The inferior occipital-frontal (IFOF) is important in connecting superior frontal and parietal cortex (Hau et al., 2016). It is impossible to determine precisely which of these tracts have the most important links with experience because of limitations of the ability of DTI to distinguish crossing fibres (Jbabdi et al., 2010), however, emerging evidence suggests that these three tracts may be reasonable candidates for future studies to explore. For example, recent work has suggested that the microstructural architecture of the SLF is predictive of patterns of unpleasant brooding in depression, and was predictive of the functional connectivity of a precuneal network of the DMN (Pisner et al., 2019). Similarly, Bonnelle and colleagues (2012) found that traumatic brain injury to a white matter path identified by probabilistic analysis had led to less efficient regulation of neural activity within the DMN by the saliency network. Notably, we recently demonstrated that the saliency network plays a critical role in the adaptive allocation of conscious attention to both internal and external foci, in part, through its relationship to both the DMN and to systems important for external attention, namely the dorsal attention network (Turnbull et al., 2019a). It would be useful, therefore, in the future to explore how the structural architecture of the brain constrains the functional activity in the cortex, and in particular the DMN, across situations varying in their reliance on internal and external modes of cognition.

