## Supplementary Table 1 for "Missing the forest because of the trees: Slower alternations during binocular rivalry are associated with lower levels of visual detail during ongoing thought"

Supplementary Materials

**Supplementary Table 1**: Multidimensional experience sampling questions, MDES

| Dimensions | Questions | 1 | 4 |
|---|---|---|---|
| Task | My thoughts were focused on the task I was performing. | Not at all | Completely |
| Future | My thoughts involved future events. | Not at all | Completely |
| Past | My thoughts involved past events. | Not at all | Completely |
| Self | My thoughts involved myself. | Not at all | Completely |
| Person | My thoughts involved other people. | Not at all | Completely |
| Emotion | The content of my thoughts was: | Negative | Positive |
| Images | My thoughts were in the form of images. | Not at all | Completely |
| Words | My thoughts were in the form of words. | Not at all | Completely |
| Vivid | My thoughts were vivid as if I was there. | Not at all | Completely |
| Detailed | My thoughts were detailed and specific. | Not at all | Completely |
| Habit | This thought had recurrent themes similar to those I have had before. | Not at all | Completely |
| Evolving | My thoughts tended to evolve in a series of steps. | Not at all | Completely |
| Deliberate | My thoughts were: | Spontaneous | Deliberate |
